# Early Insulin Signaling Abnormalities and Oxidative Stress Responses in PS19 Tau Transgenic Mice

**DOI:** 10.64898/2026.10.16.759692

**Authors:** Montasir Elahi, Irina Burd, Peixin Yang, Wei-Bin Shen

**Author notes:** Corresponding author (ME).

## Abstract

Pathological tau accumulation is a hallmark of Alzheimer’s disease (AD), yet its relationship with impaired insulin signaling remains unclear. Previously, we reported altered IRS1 phosphorylation reduced p-IRβ, and GLUT3 expression in young AD transgenic mice prior to overt pathology. Here, we investigated insulin signaling and oxidative stress pathways in the Tau P301S (PS19) mouse model. We found that alterations in phospho-IRS1, phospho-IRβ, and the glucose transporters GLUT3 and GLUT4 occur before pathogenic tau deposition. In stable tau-expressing human neuronal cell lines, tau expression was similarly associated with impaired insulin signaling. Concurrent dysregulation of insulin signaling and activation of oxidative stress-responsive kinases, accompanied by increased microglial abundance, were evident in 2- month-old PS19 mice and persisted into adulthood. Together, these findings demonstrate that insulin signaling abnormalities and oxidative stress kinase activation are early and sustained features of PS19 tauopathy, preceding the onset of overt neuropathology.

**Graphical Abstract:** 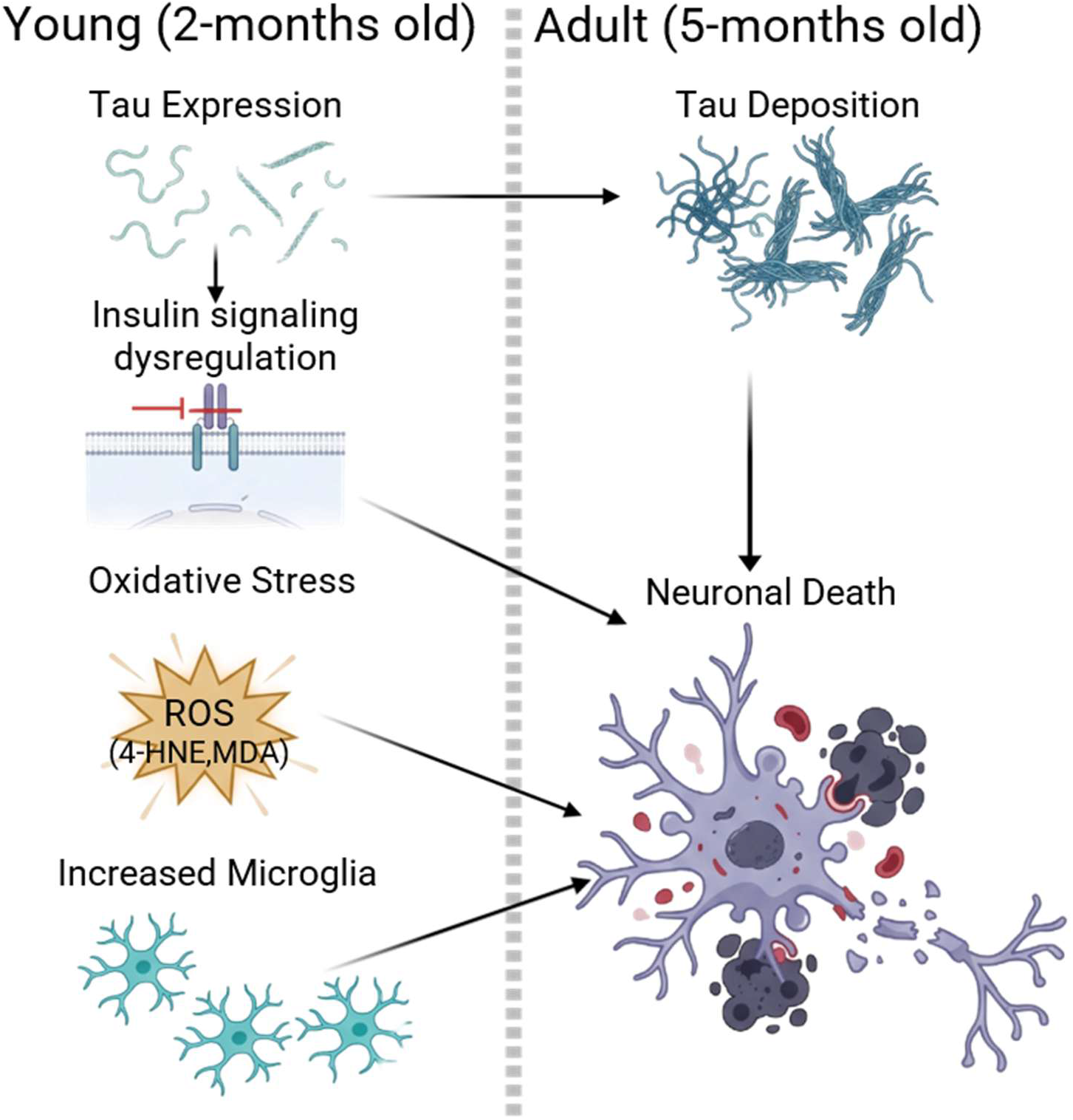

## Introduction

Alzheimer’s disease (AD) is the most common form of dementia, affecting more than 55 million people worldwide, with women accounting for nearly two-thirds of affected individuals [1, 2]. AD neuropathologically characterized by extracellular amyloid-β (Aβ) plaques and intracellular neurofibrillary tangles (NFTs), with prominent involvement of forebrain structures. Histopathological analyses have shown that neurons containing NFTs exhibit profound disruption of the cytoskeletal network, including the loss of microtubules and tubulin-associated proteins [3]. Although the molecular events linking cytoskeletal dysfunction to NFT formation are not fully understood, accumulating evidence indicates that dysregulation of protein phosphorylation and dephosphorylation pathways plays a central role in tau pathology and neurofibrillary lesion development [4]. Tau, a microtubule-associated protein and the principal component of NFTs, is extensively hyperphosphorylated in AD brains compared with age-matched non-demented controls [5]. This hyperphosphorylation occurs in a site-specific manner throughout disease progression and promotes the redistribution of tau from axons to the somatodendritic compartment. Such aberrant localization contributes to microtubule destabilization and facilitates tau aggregation and filament formation [6].

The phosphorylation state of tau is tightly controlled by multiple intracellular signaling pathways that regulate neuronal metabolism, survival, and stress responses. Among these, insulin and insulin-like growth factor-1 (IGF-1) signaling have emerged as important modulators of tau phosphorylation [7]. Tau contains more than 80 potential serine, threonine, and tyrosine phosphorylation sites [8], and accumulating evidence indicates that insulin and IGF-1 signaling influence both tau expression and phosphorylation status [9]. Consequently, disturbances in these pathways may contribute directly to tau dysregulation and neurodegeneration.

Growing evidence suggests that impaired brain insulin signaling is a major contributor to cognitive decline and neurodegeneration in AD [10, 11]. Under physiological conditions, insulin and IGF-1 regulate neuronal survival, synaptic plasticity, energy metabolism, and memory formation through activation of insulin receptors (IRs) and IGF-1 receptors (IGF-1Rs) [12]. Numerous studies have shown that IRs are abundantly expressed in the hippocampus, neocortex, hypothalamus, and cerebellum, while elevated expression of IGF-1R is exclusively observed in the hippocampus, neocortex and thalamus [13]. Activation of IR and IGF-1R initiates downstream signaling cascades such as PI3K/AKT and MAPK, which regulate metabolism, protein synthesis, cellular growth, and survival [14]. Dysregulation of these pathways has been associated with abnormal tau phosphorylation, synaptic dysfunction, and neuronal loss, supporting the concept of AD as a disorder with prominent brain insulin resistance.

In parallel, oxidative stress has emerged as another key contributor to AD pathogenesis and may represent a mechanistic link between insulin resistance and tau pathology [15, 16]. Reactive oxygen species and oxidative damage can impair insulin signaling by altering the activity of insulin receptors and downstream signaling molecules while simultaneously activating stress-responsive kinases capable of phosphorylating tau [15]. Thus, oxidative stress has the potential to amplify both metabolic dysfunction and tau pathology, creating a feed-forward cycle that promotes neurodegeneration.

There is ongoing debate as to whether oxidative stress acts as an initiating factor or a downstream consequence of AD pathology. In our recent study using post-mortem human AD brain tissues and APP/PS1 transgenic mice, we demonstrated that activation of oxidative stress-responsive kinase signaling and insulin resistance occurred before overt pathological manifestations of AD [17]. In this study, we utilized the tauopathy model mice P301S (PS19), characterized by prominent neurofibrillary tangle-like inclusions in the neocortex, hippocampus, brain stem, and spinal cord [18]. We aimed to further investigate a potential causal relationship linking alterations in insulin signaling, oxidative stress, and AD pathogenesis. Consistent with our previous studies, the present findings further support the existence of metabolic dysregulation and oxidative stress preceding the development of tau deposition related pathology.

## Materials and Methods

### Animals, sacrifice, brain tissue collection and preparation for biochemical analysis

Tau transgenic PS19 mice (P301S Tg mice) were obtained from Jackson Laboratory. These mice express the P301S mutant form of human microtubule-associated protein tau (1N4R *MAPT*) under the control of the mouse prion protein promoter (*Prnp*). PS19 mice and their non-transgenic (NTg) control littermates shared the same C57BL/6J genetic background. All animal experiments were conducted in accordance with Institutional Animal Care and Use Committee (IACUC) guidelines. Female PS19 and non-transgenic (NTg) mice aged 1, 2, and 5 months (n = 6 per group) were used for biochemical and immunohistochemical analyses. An additional four mice per group were utilized for cortical neuron isolation. Mice were euthanized by cervical dislocation, and the frontal cortex was dissected for further analyses. The left and right hemispheres were used for histological and biochemical analyses, respectively.

### Mouse brain tissue preparation for biochemical analysis and Western Blotting

Frozen frontal cortex tissues were homogenized and sonicated in a 10-fold volume of homogenization buffer (HB) containing 10 mM Tris-HCl (pH 7.5), 0.8 M NaCl, 1 mM EGTA, 1 mM dithiothreitol (DTT), and 0.1 mM phenylmethanesulfonyl fluoride (PMSF). Following centrifugation at 20,000 × g for 20 min at 4°C, the supernatants were collected and used for subsequent analyses. Protein concentrations were determined using a Pierce BCA Protein Assay Kit (Thermo Fisher Scientific), and samples were mixed with 4× loading buffer prior to western blot analysis.

Proteins were separated on 4–20% Mini-PROTEAN TGX™ Precast Gels (Bio-Rad, USA) and transferred to polyvinylidene fluoride (PVDF) membranes (Immobilon-P). To minimize non-specific binding, membranes were blocked with 5% bovine serum albumin (BSA) for 1 h and then incubated with primary antibodies overnight at 4°C. After washing, membranes were incubated with the appropriate HRP-conjugated secondary antibodies. Protein bands were visualized using Immobilon Western HRP substrate (Millipore) and enhanced chemiluminescence (ECL). Band intensities were quantified using ImageJ software (National Institutes of Health, USA).

### Immunofluorescence

Immunofluorescence staining was performed on formalin-fixed frozen sections as previously described [19]. Briefly, 10-µm-thick sagittal brain sections were rehydrated, and endogenous peroxidase activity was quenched with 0.3% H₂O₂ in methanol. Antigen retrieval was performed by autoclaving the sections at 120°C for 10 min in citrate buffer (pH 6.0). Following blocking with 10% normal goat serum (Vectastain), sections were incubated overnight at 4°C with primary antibodies, followed by incubation with species-specific Alexa Fluor-conjugated secondary antibodies. Nuclei were counterstained with 4’,6-diamidino-2-phenylindole (DAPI; Thermo Fisher Scientific). Fluorescence images were acquired using a Nikon fluorescence microscope (Eclipse Ni-E, Nikon, Japan). For each mouse, a minimum of three randomly selected images from the frontal cortex were captured per brain section.

### Neurons Isolation, RNA extraction and qPCR

Neurons were isolated from the frontal cortex of 2-month-old PS19 and NTg mice using the Adult Brain Dissociation Kit and Adult Neuron Isolation Kit (Miltenyi Biotec) according to the manufacturer’s instructions. Briefly, following brain collection, the frontal cortex was dissected on ice, minced, and enzymatically dissociated at 37°C using the brain dissociation enzyme mixture. After debris and red blood cell removal, the mixed cell suspension was incubated with an adult non-neuronal cell biotin-antibody cocktail. Non-neuronal cells were subsequently labeled with Anti-Biotin MicroBeads, and neurons were isolated using a MACS magnetic separator. The purity and viability of the isolated neuronal cells were verified according to the manufacturer’s protocol. Isolated neurons were counted and immediately subjected to RNA extraction using the Direct-zol RNA Miniprep Plus Kit (Zymo Research).

Purified RNA was reverse-transcribed using the iScript Advanced cDNA Synthesis Kit (Bio-Rad). Quantitative real-time PCR (RT-qPCR) was performed using a StepOnePlus Real-Time PCR System and SYBR Green Master Mix (IDT). Relative gene expression levels were calculated using the 2^−ΔΔCt method and normalized to the housekeeping gene *β-actin.* Primer sequences used in this study are listed in Supplementary Table 1.

### Tau stable neuronal cell line preparation, ICC and qPCR

Stable 1N4R tau without any mutation expressing neuronal cell lines were generated from human neuroblastoma SH-SY5Y cells following a previously described protocol [19]. SH-SY5Y cells were maintained in DMEM/F-12 medium (Millipore Sigma) supplemented with 10% fetal bovine serum at 37°C in a humidified incubator containing 5% CO₂. Cells were transfected with the tau expression vector (pcDNA3.1-Tau1N4R) using X-tremeGENE 9 DNA Transfection Reagent (Roche) in 6-well plates. Twenty-four hours after transfection, cells were transferred to 10-cm culture dishes containing selection medium supplemented with 300 μg/mL G418 (Geneticin; Gibco). Following one month of antibiotic selection, cells were seeded into 96-well plates using a cell sorter to obtain single-cell clones. Stable clones were subsequently screened and validated for tau expression by western blotting. Control cells were prepared using pcDNA3.1-empty vector following the same protocol. Immunocytochemistry was performed using formalin fixed, 0.3% saponin permeabilized cells in 8W chamber slides. Blocking was performed with 5.0% BSA before addition of the primary antibodies. qPCR was performed from RNA obtained from cells cultured in 6W plate at 80-90% confluency using the protocol described earlier. All cell culture experiments were performed independently five times, with each experiment considered a biological replicate (n = 5).

### Antibodies

All primary antibodies used in this study are listed in Supplementary Table 2.

### Statistical analysis

Sample distribution and variance were calculated using JMP version 12.0 software (SAS Institute, North Carolina, USA) with a confidence level of 95% and α = 0.05. Comparisons between two groups were performed using Unpair T-test followed Welch’s corrections for normally distributed data and the Mann-Whitney U test for non-normally distributed data. Statistical significance was defined as P < 0.05. Data are presented as mean ± SEM.

## Results

### Early insulin signaling abnormalities and reduced neuronal glucose transporter expression in young PS19 mice

Accumulating evidence indicates that brain insulin resistance contributes to synaptic dysfunction, impaired neuroplasticity, and cognitive decline in AD [20, 21]. Consistent with this concept, alterations in insulin receptor signaling, including impaired insulin receptor β (IRβ) activity and expression, have been reported in both AD patients and experimental AD models [22]. Reduced cortical insulin receptor levels and dysregulated receptor signaling have been proposed as key features of AD-associated metabolic dysfunction [22]. To determine whether insulin signaling is altered at an early stage of tauopathy, we examined the activated form of insulin receptor β (phospho-IRβ, Tyr972) and the inhibitory form of insulin receptor substrate-1 (phospho-IRS1, Ser636) in the frontal cortex of two-month-old PS19 mice (Fig. 1A-B). Immunofluorescence analysis revealed a marked reduction in pIRβ immunoreactivity accompanied by a significant increase in pIRS1 (Ser636) levels, affecting approximately 40% of cortical cells in PS19 mice compared with age-matched controls (Fig. 1B). These findings indicate that defects in insulin signaling are already present at 2-months old mice.

**Figure 1.**
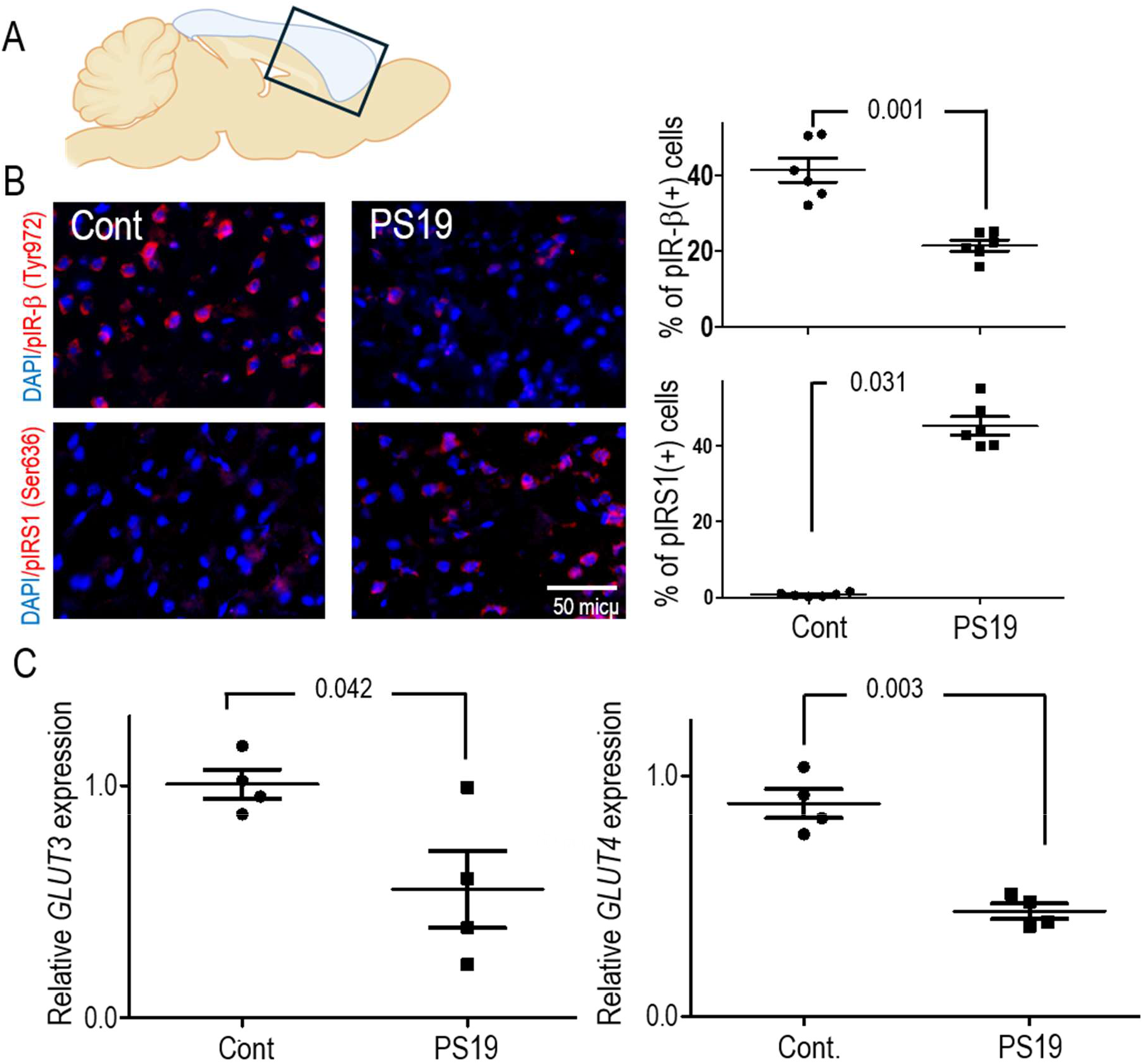
Early alterations in insulin signaling and glucose transporter expression in the frontal cortex of two-month-old PS19 mice. (A) Schematic representation of a sagittal mouse brain section indicating the frontal cortex region used for analysis. (B) Representative immunofluorescence images and quantitative analyses of phosphorylated insulin receptor β (pIRβ; Tyr972), and phosphorylated IRS1 (pIRS1; Ser636) in the frontal cortex of two-month-old PS19 and control mice (*n* = 6). Scale bars are indicated in the images. Unpair T-test followed Welch’s corrections. (C) RT-qPCR analysis of GLUT3 and GLUT4 mRNA expression in neurons isolated from the frontal cortex of two-month-old PS19 and control mice (*n* = 4). Unpair T-test followed Welch’s corrections. Data are presented as mean ± SEM. Statistical significance is indicated by respective *P* values.

Because impaired insulin signaling is closely linked to disturbances in neuronal glucose utilization, we next assessed the expression of central nervous system glucose transporters. In our previous study, we reported reduced expression of the neuronal glucose transporter GLUT3 in both AD patient brains and APP/PS1 transgenic mice [17]. To determine whether similar changes occur in a tauopathy model, we quantified the expression of glucose transporter genes in isolated frontal cortical neurons using RT-qPCR [23]. Notably, expression of the neuronal glucose transporters *GLUT3* and *GLUT4* was significantly reduced in two-month-old PS19 mice [24] (Fig. 1C). In contrast, *GLUT1* and *GLUT8* expression remained unchanged, whereas *GLUT2* and *GLUT6* transcripts were not detectable in neuron-specific RNA samples (Supplementary Fig. 1A).

To establish the temporal onset of these metabolic alterations, we further evaluated insulin resistance markers and glucose transporter expression in one-month-old PS19 mice. At this age, no significant differences were detected in insulin signaling components or glucose transporter expression, apart from elevated phosphorylation of IRS1 at Ser636 (Supplementary Fig. 2A). Together, these findings suggest that inhibitory IRS1 phosphorylation represents one of the earliest detectable metabolic abnormalities in PS19 mice and precedes the broader alterations in insulin receptor signaling and neuronal glucose transporter expression observed at two months of age.

### Early activation of oxidative stress signaling in PS19 mice prior to overt tau pathology

Oxidative stress is increasingly recognized as a central contributor to neurodegeneration, capable of inducing neuronal membrane damage and triggering apoptosis through the activation of stress-responsive kinase pathways [25]. In addition, the receptor for advanced glycation end products (RAGE) has been implicated as an important mediator of oxidative stress and inflammatory signaling in AD [26]. Consistent with this concept, our previous work demonstrated activation of oxidative stress-associated kinases, including apoptosis signal-regulating kinase 1 (ASK1), in postmortem AD brains, APOE4-associated AD-risk brains, and APP/PS1 transgenic mice [17]. To determine whether oxidative stress is an early feature of tauopathy, we examined multiple markers of oxidative damage and oxidative stress signaling in the frontal cortex of young PS19 mice. Immunofluorescence analyses revealed a clear accumulation of the lipid peroxidation products 4-hydroxynonenal (4-HNE) and malondialdehyde (MDA) in the brains of two-month-old PS19 mice, indicating the presence of early oxidative stress (Fig. 2B). This increase was accompanied by elevated expression of RAGE, further supporting the activation of oxidative stress-related pathways at this early stage of disease (Fig. 2B).

**Figure 2.**
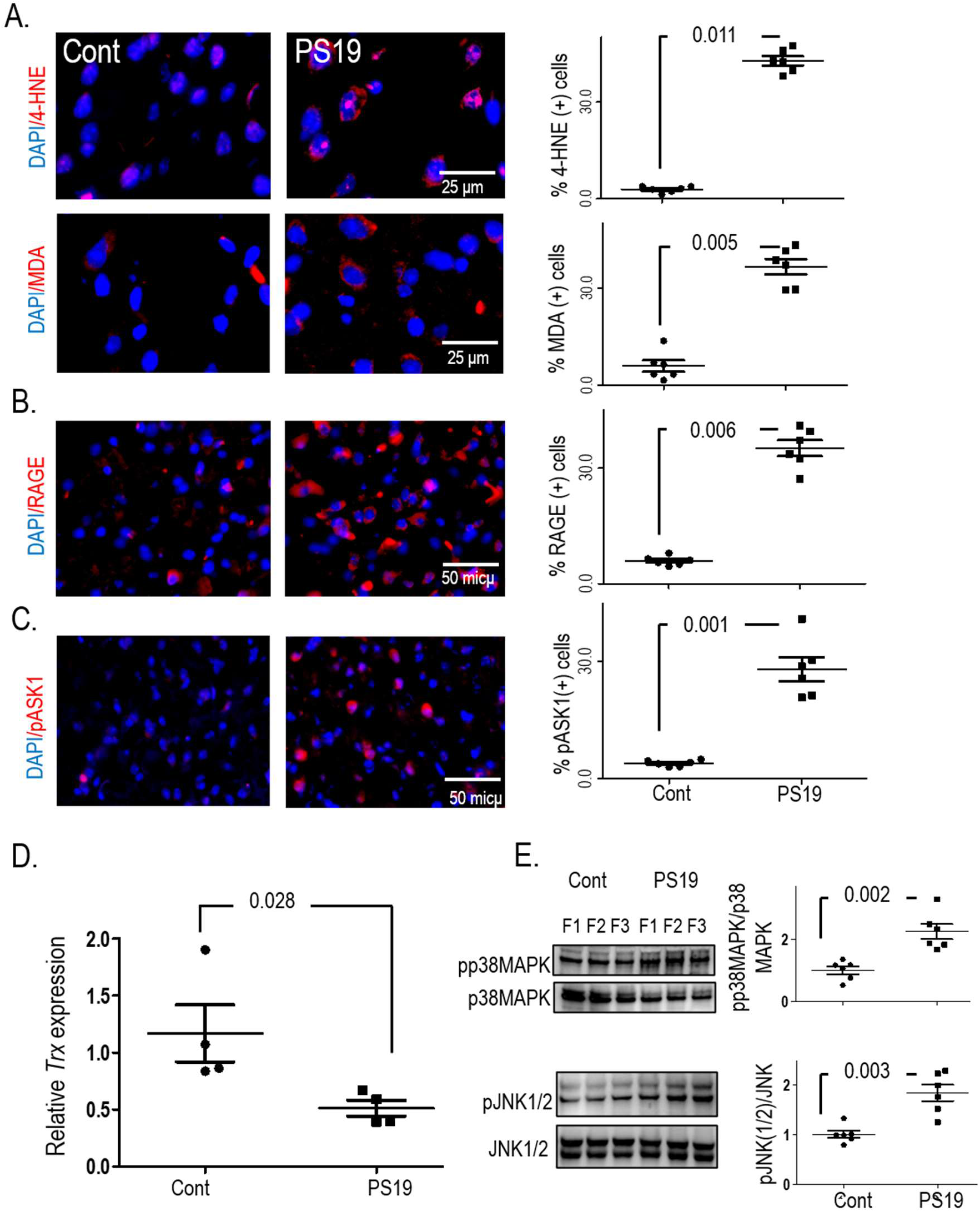
Activation of oxidative stress pathways in the frontal cortex of two-month-old PS19 mice. (A) Representative immunofluorescence images and quantitative analyses of the oxidative stress markers 4-hydroxynonenal (4-HNE) and malondialdehyde (MDA) in the frontal cortex of two-month-old PS19 and control mice. (B) Immunofluorescence analysis and quantification of receptors for advanced glycation end products (RAGE) expression in the frontal cortex of two-month-old PS19 mice. (C) Immunofluorescence analysis and quantification of activated ASK1 (phospho-Thr845-ASK1) in the frontal cortex of two-month-old PS19 mice. (*n* = 6, for all experiments) Scale bars are indicated in the respective panels. Unpair T-test followed Welch’s corrections. (D) RT-qPCR analysis of thioredoxin (Trx) mRNA expression in neurons isolated from the frontal cortex of two-month-old PS19 and control mice (*n* = 4). Mann-Whitney U test. (E) Representative western blots and quantitative analyses of phospho-p38 MAPK, and phospho-JNK1/2 expression in the frontal cortex of two-month-old PS19 and control mice (*n* = 6). Data are presented as mean ± SEM. Unpair T-test followed Welch’s corrections. Corresponding *P* values are indicated.

Given the established role of ASK1 as a key sensor and transducer of oxidative stress signals, we next evaluated its activation status. Two-month-old PS19 mice exhibited a marked increase in the activated form of ASK1, phospho-Thr845-ASK1, compared with control littermates (Fig. 2C) [27]. Activation of ASK1 has been closely linked to neurodegeneration in multiple experimental settings [28], and its activity is negatively regulated by the antioxidant protein thioredoxin [29]. Notably, neuron-specific RT-qPCR analysis demonstrated a significant reduction in thioredoxin expression in PS19 mice (Fig. 2D), suggesting a loss of endogenous antioxidant defense mechanisms that may facilitate sustained ASK1 activation. Downstream of ASK1, we observed robust activation of the stress-responsive kinases p38 MAPK and JNK1/2, as evidenced by significantly increased phosphorylation levels in the frontal cortex of two-month-old PS19 mice (Fig. 2E). Together, these findings identify a coordinated activation of oxidative stress signaling pathways characterized by oxidative damage, increased RAGE expression, reduced thioredoxin levels, and activation of the ASK1-p38/JNK signaling axis.

To define the temporal onset of these molecular changes, the same markers were examined in one-month-old PS19 mice. At this age, no significant alterations in oxidative stress markers were detected (Supplementary Fig. 2B), indicating that the oxidative stress phenotype emerges between one and two months of age, coinciding with the early metabolic and inflammatory abnormalities observed in PS19 mice.

### Early microglial activation and neuroinflammatory responses accompany tau kinase activation in PS19 mice

Microglial activation and chronic neuroinflammation are well-established features of AD pathology and are thought to contribute to disease progression [30]. Notably, activated microglia have also been reported in PS19 mice before the appearance of overt neurofibrillary tangles, suggesting that neuroinflammatory events occur early during tauopathy development [18]. To determine whether neuroinflammatory changes are present during the early stages of tauopathy, we quantified ionized calcium-binding adapter molecule 1 (Iba1)-positive microglia in the frontal cortex of two-month-old PS19 mice using immunofluorescence (Fig. 3A). As expected, Iba1-positive microglia were readily detectable in the developing brain of both PS19 and control mice. However, two-month-old PS19 mice exhibited a significant increase in the number of Iba1-positive microglia compared with age-matched littermate controls (Fig. 3A). This increase was sustained in five-month-old PS19 mice, in which microglia displayed a more pronounced activated morphology characterized by enlarged cell bodies and highly branched, spiky processes (Supplementary Fig. 3C). In contrast to microglial changes, analysis of astrocytes revealed no evidence of astrogliosis or astrocyte activation in two-month-old PS19 mice (Supplementary Fig. 1B).

**Figure 3.**
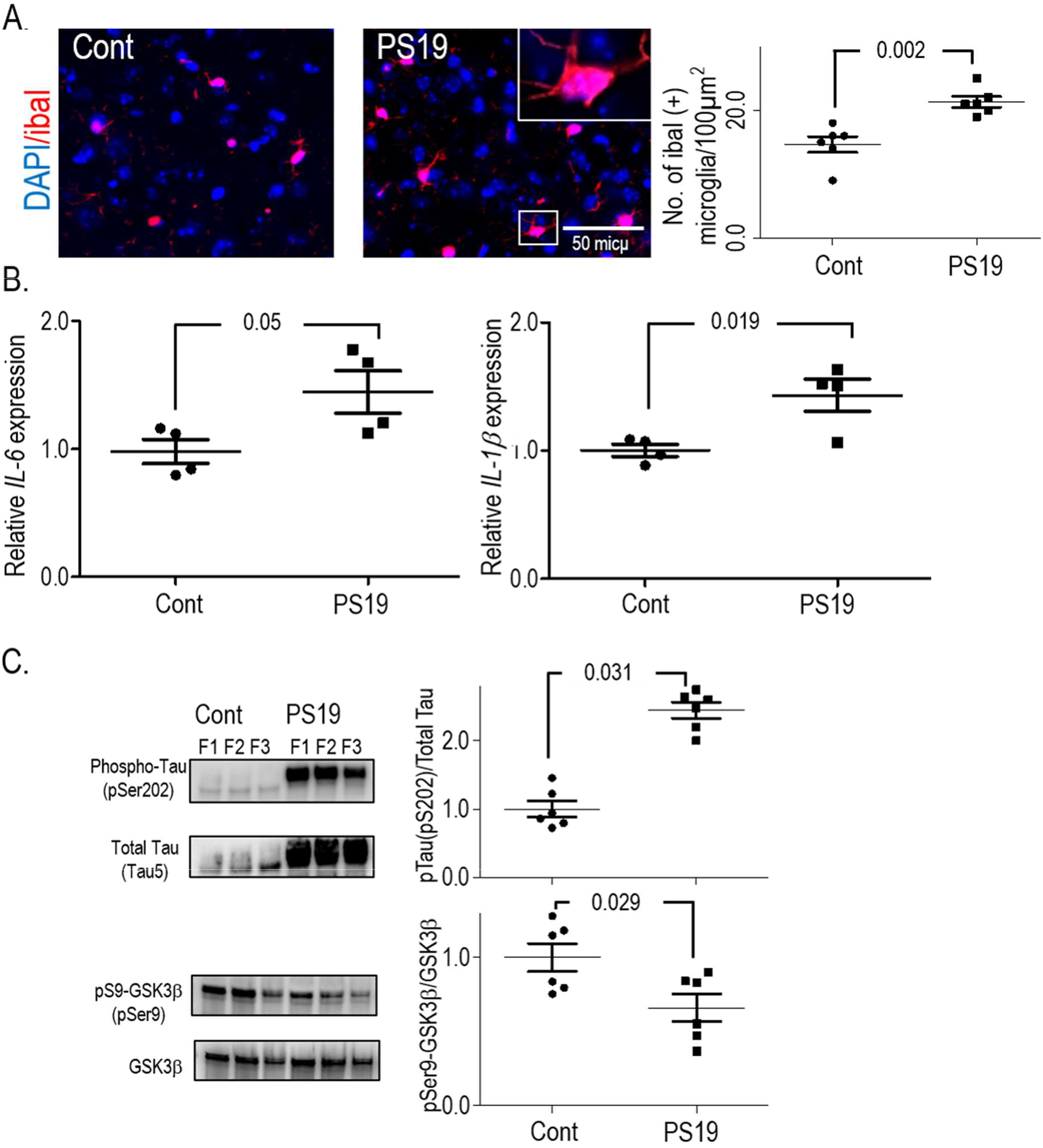
Early microglial activation, neuroinflammatory responses, and tau pathology in the frontal cortex of PS19 mice. (A) Representative immunofluorescence images and quantitative analyses of Iba1-positive microglia in the frontal cortex of two-month-old PS19 and control mice (*n* = 6). Data are presented as mean ± SEM. Unpair T-test followed Welch’s corrections. Scale bars are indicated. (B) RT-qPCR analysis of the pro-inflammatory cytokines IL-1β and IL-6 in neurons isolated from the frontal cortex of two-month-old PS19 and control mice (n=4, Unpair T-test). (C) Representative western blots and quantitative analyses of total tau & phosphorylated tau, Inhibitory form of major Tau kinase GSK-3β (pSer9 GSK 3β) & total GSK-3β in the frontal cortex of two-month-old PS19 and control mice (*n* = 6). Data are presented as mean ± SEM. Unpair T-test followed Welch’s corrections and statistical significance is indicated by the corresponding *P* values.

Consistent with enhanced microglial activation, neuronal expression of the pro-inflammatory cytokines IL-1β was elevated in PS19 mice, providing further evidence for the initiation of an early neuroinflammatory response. As IL-6 has been implicated in AD-associated cognitive dysfunction, we examined its expression in PS19 mice [31]. While IL-6 levels exhibited an upward trend compared with NTg controls, the observed increase was not statistically significant (P = 0.05) (Fig. 3B). These inflammatory changes coincided with progressive alterations in tau pathology. Increased tau phosphorylation, together with activation of the major tau kinase GSK-3β, was detected as early as one month of age and continued to increase with disease progression (Fig. 3C, Supplementary Fig. 2D). The accumulation of phosphorylated tau ultimately culminated in the formation of tangle-like structures in older animals (Supplementary Fig. 3E).

Taken together, these findings demonstrate that microglial activation, neuroinflammatory signaling, and tau kinase dysregulation emerge at very early stages of disease in PS19 mice, preceding overt tau tangle formation and supporting a potential contribution of neuroinflammation to the initiation and progression of tau pathology.

### Neuronal insulin signaling abnormalities and reduced glucose transporter expression are associated with tau expression

To investigate whether the early insulin signaling deficits observed in PS19 mice are directly linked to tau expression, we generated stable human 1N4R tau-expressing neuronal cell lines using SH-SY5Y cells. Insulin signaling was subsequently evaluated by assessing the activation status of insulin receptor β (pIRβ; Tyr972) and the inhibitory phosphorylation of IRS1 (pIRS1; Ser636). Immunofluorescence analysis revealed a significant reduction in pIRβ immunoreactivity accompanied by a marked increase in pIRS1 (Ser636) levels in tau-expressing cells compared with control cells (Fig. 4A). These findings closely mirror the alterations observed in young PS19 mice and suggest that tau expression itself is sufficient to disrupt key components of the insulin signaling cascade.

**Figure 4.**
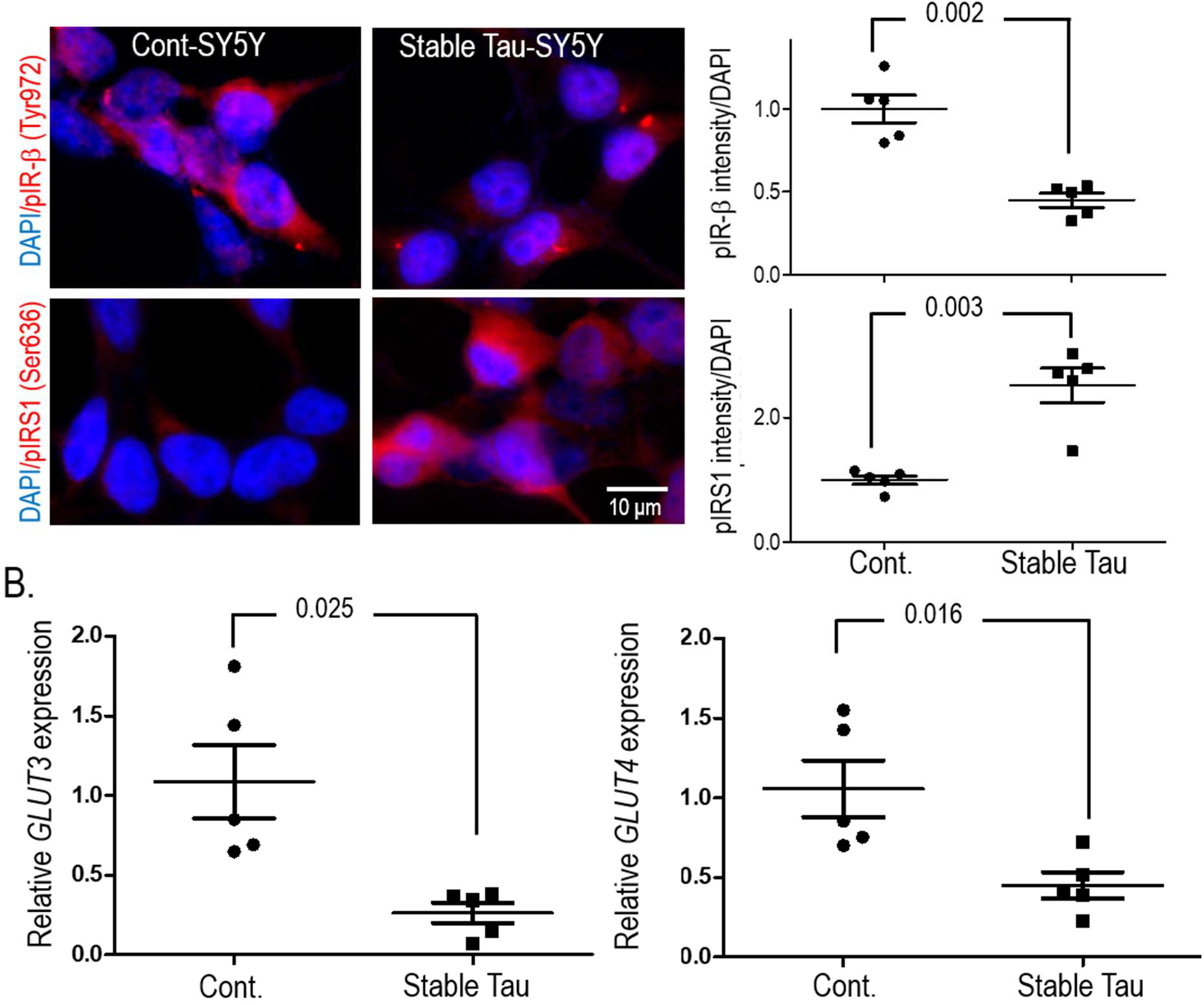
Impaired insulin signaling in stable human 1N4R tau-expressing neuronal cell lines. (A) Representative immunofluorescence images and quantitative analyses of phosphorylated insulin receptor β (pIRβ; Tyr972), and phosphorylated IRS1 (pIRS1; Ser636), in stable human 1N4R tau-expressing neuronal cells and control cells (n = 5 independent experiments). Scale bars are indicated. (B) RT-qPCR analysis of GLUT3 and GLUT4 mRNA expression in stable human 1N4R tau-expressing neuronal cells and control cells (n = 5 independent experiments). Data are presented as mean ± SEM, Unpair T-test followed Welch’s corrections. Corresponding P values are indicated.

Given the close relationship between insulin signaling and neuronal glucose utilization, we next examined the expression of the neuronal glucose transporters GLUT3 and GLUT4. Consistent with our in vivo observations, both transporters were significantly downregulated in stable tau-expressing cells relative to control cells (Fig. 4B). The concordance between the cellular and animal models further supports a direct association between tau expression, impaired insulin signaling, and reduced neuronal glucose transport capacity.

Remarkably, despite the robust alterations in insulin signaling and glucose transporter expression, no significant changes were detected in oxidative stress markers in 1N4R tau-expressing cells (Supplemental Fig. 4). These findings suggest that disturbances in insulin signaling may arise as an early consequence of tau expression, whereas the induction of oxidative stress may require additional pathogenic properties associated with disease-linked tau mutations.

## Discussion

A defined molecular link between Alzheimer’s disease (AD) and diabetes has yet to be established. Epidemiological and experimental studies consistently identify diabetes as a major risk factor for sporadic AD, largely through its association with insulin resistance, dyslipidemia, oxidative stress, and chronic inflammation, all of which are increasingly recognized as fundamental contributors to neurodegeneration [32]. Several studies have further demonstrated that these pathological processes promote the expression and activation of the receptor for advanced glycation end products (RAGE), a key mediator of oxidative stress and inflammatory signaling that is strongly associated with AD pathology [33, 34]. In our previous study, we reported early impairment of insulin signaling accompanied by activation of oxidative stress pathways, ASK1 signaling, and downstream stress kinases in both young APP/PS1 mice and APOE4-associated AD-risk brains [17]. The present study extends these observations to a highly reported tauopathy model and demonstrates that abnormalities in insulin signaling, oxidative stress pathways, neuroinflammatory responses, and neuronal glucose transport emerge long before overt tau deposition in PS19 mice.

One of the most notable findings of the present study is the appearance of insulin signaling abnormalities at a remarkably early stage of disease progression. Brain insulin resistance has emerged as a defining metabolic feature of AD, supported by extensive evidence demonstrating dysregulation of insulin receptors (IRs), insulin receptor substrates (IRSs), and insulin-like growth factor signaling pathways in vulnerable brain regions [13, 35]. Recent studies have further shown that cerebrospinal fluid levels of the soluble insulin receptor correlate with Aβ42/Aβ40 ratios, highlighting a close association between impaired insulin signaling and AD pathology [36]. Likewise, increased inhibitory phosphorylation of IRS1 at Ser636/639 has been strongly associated with amyloid burden and disease severity in postmortem AD brains [21]. Consistent with these observations, we detected a reduction in the activated form of IRβ (pIRβ-Tyr972) together with increased inhibitory phosphorylation of IRS1 (pIRS1-Ser636) in the frontal cortex of two-month-old PS19 mice. Importantly, these molecular alterations occurred long before the appearance of overt neurofibrillary tangle-like pathology, which becomes prominent in PS19 mice at later stages of disease (Supplementary Fig. 3D). These findings suggest that insulin resistance is not merely a downstream consequence of neurodegeneration but may represent an initiating pathological event in tau-mediated disease.

To further investigate whether tau itself contributes to insulin signaling dysfunction, we generated stable human 1N4R tau-expressing neuronal cell lines. Remarkably, expression of tau alone was sufficient to reproduce the insulin signaling defects observed in vivo, including reduced pIRβ and increased pIRS1 (Ser636). These findings provide direct evidence that tau expression can disrupt neuronal insulin signaling independently of advanced pathology or associated cellular changes occurring in the brain [16]. Several studies have reported reciprocal interactions between tau biology and insulin signaling, whereby impaired insulin signaling promotes abnormal tau phosphorylation, while pathological tau further exacerbates metabolic dysfunction through disruption of insulin receptor trafficking and downstream signaling cascades [9, 37]. Our findings support this model and suggest that tau-induced metabolic dysfunction may represent an early event capable of initiating a self-reinforcing pathogenic cycle.

Defective insulin signaling is closely linked to disturbances in cerebral glucose metabolism, another hallmark of AD. Brain glucose hypometabolism is detectable years before clinical symptom onset and can be visualized by ^18^F-FDG positron emission tomography in both AD patients and individuals at high risk of developing the disease [38, 39]. Consistent with impaired neuronal glucose utilization, altered expression of *GLUT3/4* has been reported in both human AD brains and transgenic mouse models [40]. In our previous work, reduced *GLUT3* expression was observed in pre-symptomatic APP/PS1 mice and AD carrier *ApoE4/4* brains [17]. In the current study, neuronal *GLUT3* and *GLUT4* expression were significantly reduced in two-month-old PS19 mice. Notably, identical reductions were observed in stable tau-expressing neuronal cells, supporting a direct relationship between tau expression and impaired glucose transport. Given that *GLUT3* serves as the principal neuronal glucose transporter and *GLUT4* contributes to activity-dependent glucose uptake, simultaneous reductions in both transporters may substantially compromise neuronal bioenergetics and synaptic function [41, 42]. This early Insulin signaling abnormality in mice comparable to insulin resistance-associated mild cognitive impairment (MCI) in human [43] and aligns with our previous report demonstrating insulin resistance in AD risk factor *ApoE4/4* carriers [17].

A second major finding of this study is the early activation of oxidative stress pathways. Oxidative stress has long been implicated in both peripheral insulin resistance and AD pathogenesis. In peripheral tissues, excess reactive oxygen species (ROS) contribute directly to insulin resistance by disrupting insulin receptor signaling [44]. In the central nervous system, however, ROS generation appears to have a more complex role and may initially arise as a compensatory response to metabolic dysfunction and altered neuronal signaling [45]. Notably, we detected no significant increase in oxidative stress markers in one-month-old PS19 mice. By two months of age, however, substantial accumulation of the lipid peroxidation products 4-HNE and MDA was evident, indicating the onset of measurable oxidative damage. This temporal pattern is particularly informative because tau hyperphosphorylation was already detectable before overt oxidative stress became apparent. Such findings suggest that early metabolic disturbances may initially trigger adaptive stress responses that subsequently progress to sustained oxidative injury as pathology advances.

The concomitant increase in RAGE expression further supports the involvement of a ROS-RAGE signaling axis during the early stages of tauopathy. RAGE is activated by a diverse range of ligands, including advanced glycation end products, oxidized lipids, amyloid-β, and damage-associated molecular signals, and acts as a potent amplifier of oxidative stress and inflammation [46]. Sustained activation of RAGE promotes ROS production through NADPH oxidase activation and establishes a feed-forward cycle that perpetuates cellular stress and neuroinflammation [46, 47]. Consequently, elevated RAGE expression in young PS19 mice may represent an important mechanistic link connecting metabolic dysfunction, oxidative stress, and progressive tau pathology [24, 48].

Consistent with increased oxidative stress, we observed robust activation of ASK1 and its downstream signaling pathways. ASK1 functions as a redox-sensitive kinase that is normally suppressed by thioredoxin under physiological conditions [49]. Oxidative stress disrupts the ASK1-thioredoxin complex, resulting in ASK1 activation and subsequent stimulation of the p38 MAPK and JNK pathways. In agreement with this mechanism, PS19 mice exhibited reduced thioredoxin expression, increased phospho-ASK1, and activation of both p38 MAPK and JNK. ASK1-mediated activation of p38 MAPK has previously been implicated in tau phosphorylation and neurodegeneration, and increased formation of ASK1-p38MAPK-tau complexes has been reported in AD brains [50]. Similarly, numerous studies have established a critical role for JNK signaling in AD pathogenesis, demonstrating that modulation of JNK activity alters amyloid burden, tau phosphorylation, and neuronal survival [51–53]. Together, our findings suggest that oxidative stress-dependent activation of the ASK1-p38/JNK axis may represent an important molecular mechanism linking early insulin resistance to progressive tau pathology.

Another notable observation was the emergence of neuroinflammatory changes shortly after the development of insulin resistance and oxidative stress signaling. Activated microglia have been reported in AD brains and are present in PS19 mice before overt tangle formation [18, 30, 54]. Consistent with these reports, we observed increased numbers of activated Iba1-positive microglia together with elevated neuronal expressions of IL-1β and IL-6. These findings suggest that neuroinflammatory processes are initiated during the earliest stages of tauopathy and likely interact with metabolic and oxidative stress pathways. Indeed, increasing evidence indicates that insulin resistance, oxidative stress, and neuroinflammation do not occur independently but instead form an interconnected pathological network in which each process amplifies the others, ultimately accelerating neuronal dysfunction and degeneration.

Collectively, our findings support a model in which tau-associated metabolic dysfunction emerges before overt neuropathology and initiates a cascade of interconnected events involving insulin signaling impairment, reduced neuronal glucose transport, oxidative stress, activation of the ASK1-p38/JNK pathway, and neuroinflammatory responses (Fig. 5). Importantly, the observation that stable tau expression alone is sufficient to induce insulin signaling abnormalities and reduce GLUT3 and GLUT4 expression suggests that metabolic dysfunction may occur as a direct consequence of tau expression rather than solely as a secondary response to established pathology. These results provide further evidence that alterations in neuronal metabolism represent early and persistent features of tauopathy and reinforce the growing view that insulin signaling pathways constitute promising therapeutic targets for AD intervention.

**Figure 5.**
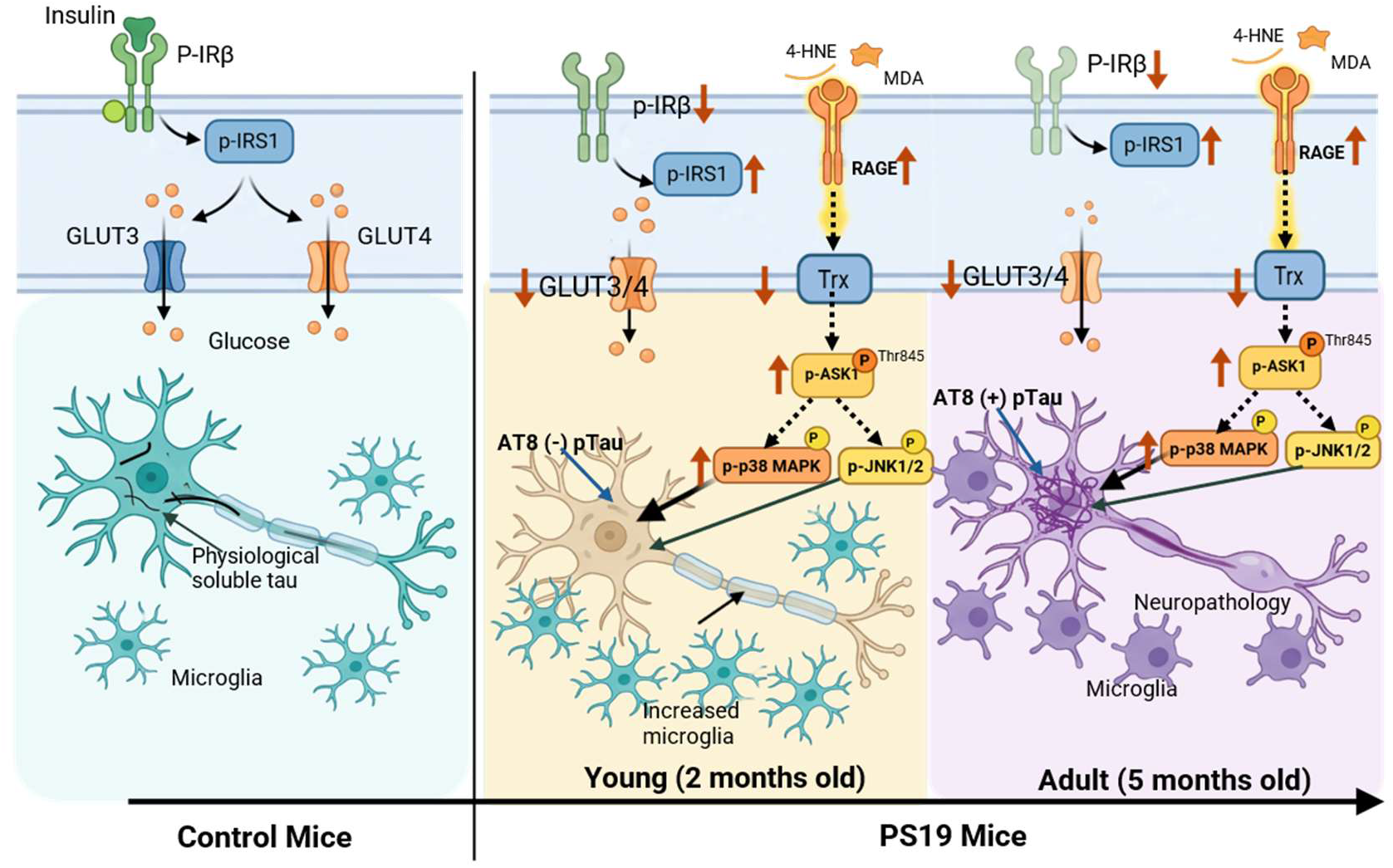
Proposed model summarizing the study findings. Tau overexpression in PS19 mice induced early disruption of insulin signaling, characterized by reduced IR-β phosphorylation (Tyr972), increased inhibitory IRS-1 phosphorylation (Ser636), and decreased expression of the neuronal glucose transporters GLUT3 and GLUT4. Notably, these alterations were evident as early as 2 months of age and persisted through adulthood (5 months). Concurrently, markers of oxidative stress, including 4-HNE, MDA, and RAGE, were significantly elevated, accompanied by activation of oxidative stress-responsive kinases. Importantly, these molecular abnormalities emerged well before the appearance of detectable AT8-positive tau in the frontal cortex, suggesting that insulin signaling dysregulation and oxidative stress are among the earliest pathogenic events preceding overt tau deposition in the PS19 brain.

## Conclusion

The present study demonstrates that insulin signaling abnormalities, reduced glucose transporter expression, oxidative stress pathway activation, and neuroinflammatory responses arise well before overt tau deposition in PS19 mice. The replication of key metabolic defects in stable tau-expressing neuronal cells further supports a direct role for tau in initiating these pathological changes. Together with our previous findings in APP/PS1 mice and APOE4-associated AD-risk brains, these data suggest that metabolic dysfunction represents a convergent and early event in AD pathogenesis. Targeting insulin signaling and associated metabolic pathways may therefore offer a valuable strategy for therapeutic intervention during the preclinical stages of tau-mediated neurodegeneration.

## Acknowledgments

Authors are thankful to Dr. Mahadi Hassan for his occasional help with tissue sectioning.

## Contribution

M.E. Concept and designed research, performed experiments, and analyzed data; W. S. helped with the experiments and data analysis. P.Y. supervised the study. P.Y. and I. B. contributed new reagents/analytic tools; M.E. wrote the manuscript.

## Availability of the data, methods and materials

All Western blotting and immunofluorescent data will be deposited in public domains and detailed methods and reagents will be provided upon request.

