## Supplemental File for "Early Insulin Signaling Abnormalities and Oxidative Stress Responses in PS19 Tau Transgenic Mice"

**Supplemental Table 1: Detail about the qPCR primers used in this study**

| Gene name | direction | Sequence (Mouse) | Sequence (Human) |
|---|---|---|---|
| *Glut1* | F | TCAACACGGCCTTCACTG | TTGCAGGCTTCTCCAACTGGAC |
| | R | CACGATGCTCAGATAGGACATC | CAGAACCAGGAGCACAGTGAAG |
| *Glut2* | F | TGTGCTGCTGGATAAATTCGCCTG | ATGTCAGTGGGACTTGTGCTGC |
| | R | AACCATGAACCAAGGGATTGGACC | AACTCAGCCACCATGAACCAGG |
| *Glut3* | F | TTCTGGTCGGAATGCTCTTC | TGCCTTTGGCACTCTCAACCAG |
| | R | AATGTCCTCGAAAGTCCTGC | GCCATAGCTCTTCAGACCCAAG |
| *Glut4* | F | GTAACTTCATTGTCGGCATGG | CCATCCTGATGACTGTGGCTCT |
| | R | AGCTGAGATCTGGTCAAACG | GCCACGATGAACCAAGGAATGG |
| *Glut5* | F | GGCTCATCTTCCCCTTCATTC | GGCTTCTCCATCTGCCTCATAG |
| | R | ATGAATGTCCTGCCCTTGG | GGAGATGACACAGACGATGCTG |
| *Glut6* | F | TTGGTGCTGTGAGGCT | CCGGACTACGACACCTTCC |
| | R | TGGCACAAACTGGACGTA | GGATGTGTAGACCAGGGCATA |
| *Glut8* | F | TTCATGGCCTTTCTAGTGACC | GTCTCTGCACAGCCTGTTGATG |
| | R | GAGTCCTGCCTTTAGTCTCAG | GATCTCTGACATGAGGAGCCAG |
| *Txn* | F | CCCACTTGCCCCAACTGTT | GTAGTTGACTTCTCAGCCACGTG |
| | R | GGGAGTGTCTTGGAGGGAC | CTGACAGTCATCCACATCTACTTC |
| *IL 1β* | F | GCAACTGTTCCTGAACTCAACT | ATGATGGCTTATTACAGTGGCAA |
| | R | TTGGTCCTTAGCCACTCCTTC | GTCGGAGATTCGTAGCTGGA |
| *IL 6* | F | TAGTCCTTCCTACCCCAATTTCC | ACTCACCTCTTCAGAACGAATTG |
| | R | ATCTTTTGGGGTCCGTCAACT | CCATCTTTGGAAGGTTCAGGTTG |
| *Actin* | F | CATTGCTGACAGGATGCAGAAGG | CATGTACGTTGCTATCCAGGC |
| | R | TGCTGGAAGGTGGACAGTGAGG | CTCCTTAATGTCACGCACGAT |

**Supplemental Table 2: Detail about the antibodies used in this study**

| Target | Clones | Source | Application | Dilution |
|---|---|---|---|---|
| Total Tau (aa 210-241) | Tau5 | MilliporeSigma #MAB361 | Western blotting Immunofluorescence | 1:1000, ECL 1:1000 |
| phospho-Tau (pSer202) | Polyclonal | In-house | Western blotting | 1:2000, ECL |
| phospho-Tau (pSer202?Thr205)-AT8 | Mouse-monoclonal | Invitrogen#MN1020 | | |
| phospho-IRβ (Tyr 972) | Rabbit-mono | Invitrogen#44800G | Immunofluorescence , Western blotting | 1:200, 1:1000 ECL |
| phospho-IRβ (Tyr 972) | Polyclonal | Abcam#ab5678 | Immunofluorescence | 1:100 |
| Insulin receptor-β | Rabbit-mono | CST (4B8) #3025 | Western blotting | 1:1000 ECL |
| phospho-IRS-1 (pSer636) | Polyclonal | Abcam# ab277770 | Immunofluorescence | 1:50 |
| phospho-IRS-1 (pSer636) | Polyclonal | CST #2388 (AB_330339) | Immunofluorescence, Western blotting | 1:50, 1:1000 ECL |
| IRS-1 | Polyclonal | CST #2382 (AB_330333) | Western blotting | 1:1000, ECL |
| Glut3 | Polyclonal | Protech#20403-1-AP | Immunofluorescence | 1:200 |
| Glut4 | Rabbit-mono | abcam#ab33780 | Immunofluorescence | 1:500 |
| Thioredoxin | Polyclonal | CST#2298 | Western blotting | 1:1000, ECL |
| RAGE | Goat-Polyclonal | R&D#AF1145 | Immunofluorescence | 1;100 |
| 4-hydroxy nonenal (4-HNE) | polyclonal | Bioss#bs-6313r | Immunofluorescence | 1:500 |
| Melondealdehyde (MDA) | polyclonal | Genetex#GTX12835 | Western blotting | 1:1000 ECL |
| pASK1(Thr845) | polyclonal | Invitrogen#PA5-99389 | Immunofluorescence | 1:50 |
| ASK1 | polyclonal | CST##8662 | Western blotting | 1:1000, ECL |
| phospho-p38MAPK (pThr180/pTyr182) | Polyclonal | CST #9211(AB_331641) | Western blotting | 1:1000, ECL |
| p38 MAPK | Polyclonal | CST #9212(AB_330713) | Western blotting | 1:1000, ECL |
| phospho-JNK (pThr183/pTyr185) | Polyclonal | CST #9251(AB_331659) | Western blotting | 1:1000, ECL |
| JNK | Polyclonal | CST #9252 (AB_2250373) | Western blotting | 1:1000, ECL |
| Iba I | Polyclonal | WAKO#019-19741 | Immunofluorescence | 1:1000 |
| GFAP (2E1E8) | Mouse-monoclonal | Biolegends#644702 | Immunofluorescence | 1:50 |
| GAPDH | GAPDH-71.1 | Millipore Sigma#G8795 | Western blotting | 1:1000, ECL |
| Actin | AC-15 | Millipore Sigma#A5441 | Western blotting | 1:1000, ECL |

**S. Figure Legends:**

**Supplementary Fig. 1: Analysis of the insulin signaling two -month-old PS19 mice brain frontal cortex. (A)** Quantitative real time PCR analysis of *GLUT1* and *GLUT8* gene expression in the frontal cortex of two-month-old PS19 mice (n = 4)**.** Unpair T-test followed Welch's corrections. **(B)** Representative **i**mmunofluorescence images of Glial fibrillary acidic protein (GFAP) in the frontal cortex of two-month-old PS19 mice (n = 6). Scale bar is indicated.

**Supplementary Fig. 2: Analysis of the insulin signaling, oxidative stress and tau in one -month-old PS19 mice brain frontal cortex. (A)** Immunofluorescence images (Upper 3 panels) and quantification analysis (lower panel) of the phospho-Insulin receptor-β (Tyr 972) (first panel) and phosphor-IRS1 (Ser636) (second panel) in the frontal cortex of one-month-old PS19 mice (n = 6). Unpair T-test followed Welch's corrections and Scale bar is indicated. **(B)** Immunofluorescence analysis of the oxidative stress markers 4-HNE and MDA, and the stress kinase ASK1 in frontal cortex of one-month-old PS19 mice (n=6, Unpair T-test followed Welch's corrections). Scale bars are presented accordingly. **(C)** Western blotting and quantification analysis of Thioredoxin and phosphorylation of p38MAPK, JNK1/2 and Tau in frontal cortex of one-month-old PS19 female mice (n=6, Unpair T-test followed Welch's corrections). **(D)** Western blots and quantitative analyses of total tau & phosphorylated tau, Inhibitory form of major Tau kinase GSK-3β (pSer9 GSK 3β) & total GSK-3β in the frontal cortex of one-month-old PS19 and control mice (n=6, Unpair T-test followed Welch's corrections). Significant p values are presented.

**Supplementary Fig. 3: Analysis of the insulin signaling oxidative stress, tau phosphorylation in five -month-old PS19 mice brain frontal cortex. (A)** Immunofluorescence images (upper 2 panels) and quantification analysis (lower panel) of the phospho-Insulin receptor-β (Tyr 972) (first panel), phosphor-IRS1 (Ser636) (second panel) in the frontal cortex of five-month-old PS19 mice (mice (n=6, Unpair T-test followed Welch's corrections, Significant p-value presented). **(B)** Immunofluorescence images analysis of the oxidative stress markers 4-HNE (first panel), the stress kinase ASK1 (second panel) and quantification (lower panel) in the frontal cortex of five-month-old PS19 mice (mice (n=6, Unpair T-test followed Welch's corrections). Scale bars are presented accordingly. **(C)** Representative immunofluorescence images of Iba1-positive microglia in the frontal cortex of five-month-old PS19 and control mice (n=6). Scale bars are indicated. **(D)** Western blotting and quantification analysis of Trx, p38MAPK and JNK1/2 phosphorylation mice (n=6, Unpair T-test followed Welch's corrections, Significant p-values are presented) (**E)** Immunofluorescence of the AT8 positive tau deposition in frontal cortex of one-, two-and five -month-old PS19 mice. Scale bar is indicated.

**Supplementary Fig. 4. Analysis of the oxidative stress in stable human 1N4R tau-expressing neuronal cell lines. (A)** Representative immunofluorescence images of 4-HNE (upper panel) and MDA (lower panel), in stable human 1N4R tau-expressing neuronal cells and control cells (n = 5). Scale bars are indicated. **(B)** RT-qPCR analysis of *Tau* mRNA expression in stable human 1N4R tau-expressing neuronal cells and control SHSY5Y cells (n = 5). Data were analyzed by Unpair T-test followed Welch's corrections; Significant p-values are presented.

**Supplementary. Fig-1**

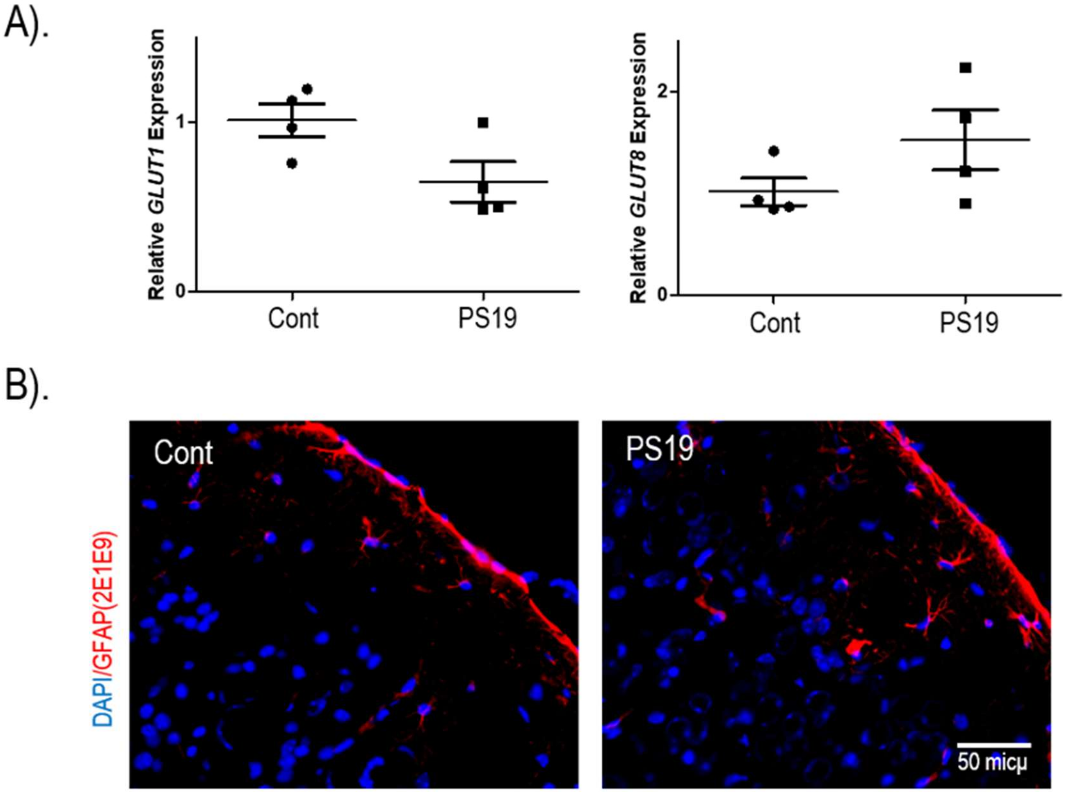

**Supplementary Fig-2**

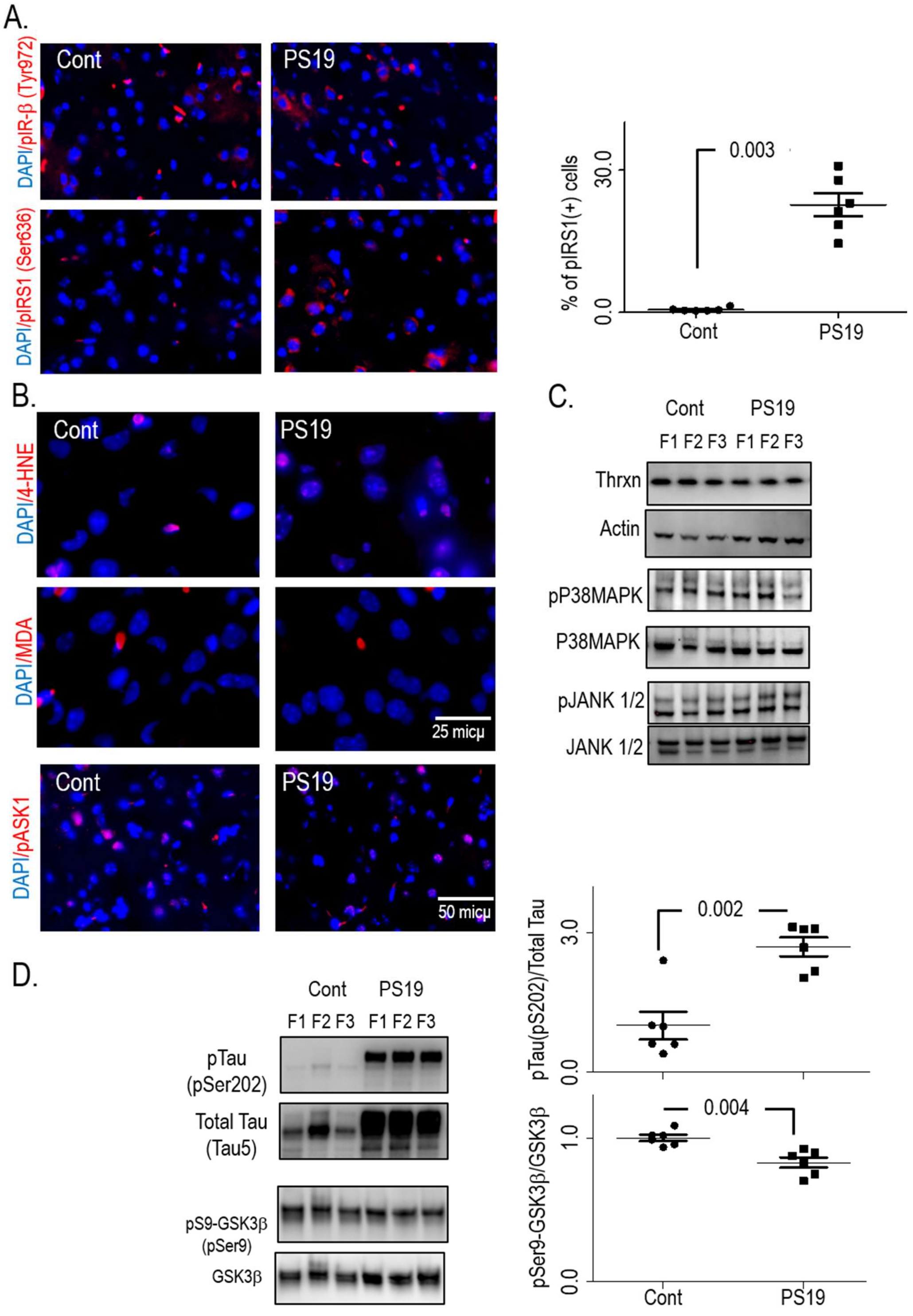

**Supplementary Fig-3**

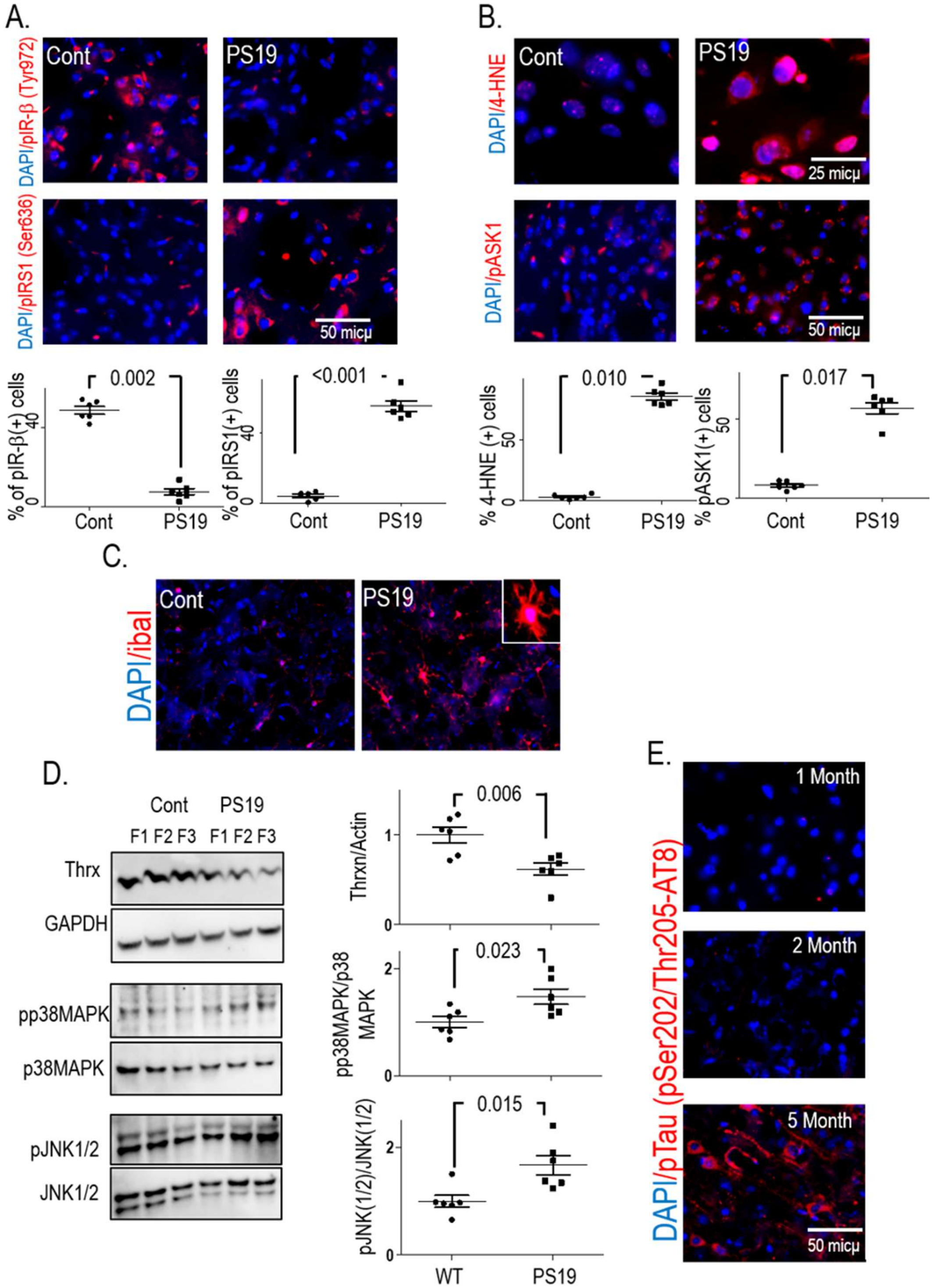

**Supplementary Fig-4**

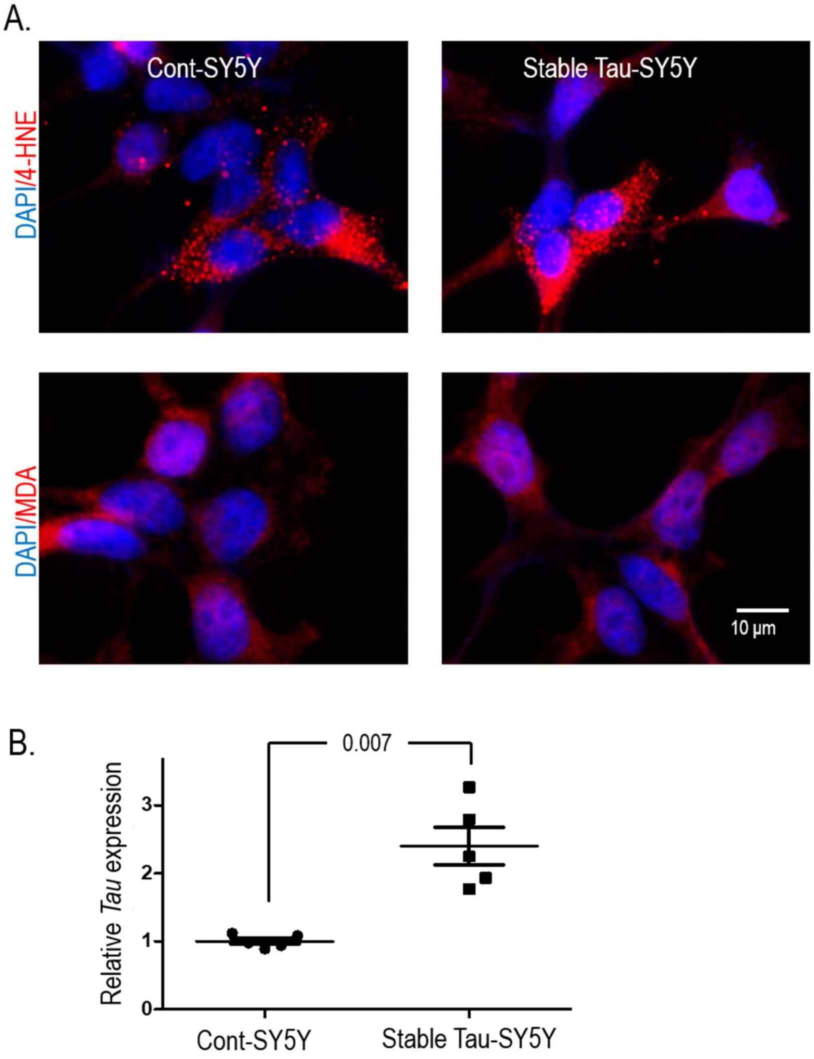
